# Language Models for the Prediction of SARS-CoV-2 Inhibitors

**DOI:** 10.1101/2021.12.10.471928

**Authors:** Andrew E Blanchard, John Gounley, Debsindhu Bhowmik, Mayanka Chandra Shekar, Isaac Lyngaas, Shang Gao, Junqi Yin, Aristeidis Tsaris, Feiyi Wang, Jens Glaser

## Abstract

The COVID-19 pandemic highlights the need for computational tools to automate and accelerate drug design for novel protein targets. We leverage deep learning language models to generate and score drug candidates based on predicted protein binding affinity. We pre-trained a deep learning language model (BERT) on ∼9.6 billion molecules and achieved peak performance of 603 petaflops in mixed precision. Our work reduces pre-training time from days to hours, compared to previous efforts with this architecture, while also increasing the dataset size by nearly an order of magnitude. For scoring, we fine-tuned the language model using an assembled set of thousands of protein targets with binding affinity data and searched for inhibitors of specific protein targets, SARS-CoV-2 Mpro and PLpro. We utilized a genetic algorithm approach for finding optimal candidates using the generation and scoring capabilities of the language model. Our generalizable models accelerate the identification of inhibitors for emerging therapeutic targets.

## 1 JUSTIFICATION FOR PRIZE

We:

- pre-train a BERT model on a dataset of 9.6 billion molecules, nearly an order of magnitude larger than previous efforts (1.1-1.6 billion) [1, 2],
- achieve 603 petaflops in mixed precision on 4032 Summit nodes, reducing pre-training time-to-solution from days to hours, and
- train a general model for protein binding affinity, accelerating the search for drug candidates relevant to SARS-CoV-2.

## 2 PERFORMANCE ATTRIBUTES

| Performance attribute | Our submission |
| --- | --- |
| **Category of achievement** | Time to solution, scalability |
| **Type of method used** | Machine learning |
| **Results reported for** | Whole application with and without I/O |
| **Precision reported** | Mixed precision (FP16 and FP32) |
| **System scale** | Measured on full-scale system (Summit) |
| **Measurement mechanism** | Internal timers, DeepSpeed FLOPS profiler |

## 3 OVERVIEW OF THE PROBLEM

The COVID-19 pandemic has drastically altered living conditions in countries throughout the world over the past two years. To date, approximately 230 million people have been infected and 4.7 million have been killed by variants of the SARS-CoV-2 virus [3]. It is not unrealistic to assume that another event like this is possible; several infectious diseases with the potential for global impact have been documented in recent years, including SARS, MERS, Ebola, and Zika [4]. Within this broader context, the current pandemic highlights the need for the development of therapeutic agents to combat emerging infectious diseases. Unfortunately, the speed at which antivirals have been developed has not maintained pace with the frequency of outbreaks. For example, although vaccines have been developed as an effective means to prevent SARS-CoV-2 infection, no clinically tested therapeutics have been approved for widespread use except for antibody treatment [5–7]. Furthermore, recent clinical trials highlight the continued need for antivirals [8]. Therefore, the timely development of drugs to treat emerging viral threats, in combination with preventive vaccines, poses a key challenge with global implications.

Although many previous efforts in drug discovery have been successful, the process can be prohibitively long (i.e., 10 to 15 years) for response to an emerging pandemic. The approval of a single compound for widespread use typically involves the screening of small molecules for potential candidates, hit-to-lead (H2L) testing followed by extensive multi-stage clinical trials [9]. The initial step of determining interesting molecules for further investigation is pivotal due to the vast size of chemical space, which prevents an exhaustive search using costly experiments and trials. To accelerate the screening process, tools from machine learning (ML) and high-performance computing (HPC) have been increasingly used to guide the selection of promising drug candidates [10–12]. Although computational methods can partially alleviate some of the associated experimental costs, typical approaches require the creation of a large compound library with measured properties for ML model training [11, 12]. Therefore, a timely response to an emerging pandemic also poses a challenge for computational methods, as custom models and datasets must be quickly generated for the new targets of interest.

To overcome the challenges associated with accelerating the discovery of drug candidates for novel protein targets, a computational approach is needed that satisfies the following criteria: (i) leverages existing large compound libraries without the need for chemical property measurements; (ii) predicts affinities for novel protein targets with limited or no additional experimental data; (iii) explores chemical space to efficiently identify compounds for further investigation. To satisfy the three criteria, we leverage high performance computing (HPC) to train generalizable ML models for both candidate generation and affinity prediction.

To take advantage of large existing compound libraries, we utilize a text representation for molecule data known as SMILES, Simplified Molecular Input Line Entry System [13]. Using Enamine *REAL* database [14] as a starting point, we generate a novel dataset of approximately 9.6 billion unique molecules. The dataset is used to pre-train a Transformer model (i.e. BERT), using the mask prediction task commonly found in natural language processing applications. During pre-training, sub-sequences of a given molecule are replaced by a mask, and the model must predict the appropriate sequence based on context. Therefore, the model learns a representation for chemical structure in a completely unsupervised manner that does not require additional property measurements.

To predict affinities for protein targets, we fine-tune the pretrained molecule model on a dataset with over a million known protein and ligand binding affinities. The fine-tuned model utilizes two pre-trained language models to generate embeddings for a given molecule and protein. For the protein embedding, we utilize a recently published Transformer model for protein sequences [15]. By using models for molecules and proteins trained in an unsupervised manner on large datasets, the fine-tuned model leverages the structural information in the respective embeddings. A final cross attention layer is added on top of the embeddings to generate an affinity score for any given protein and molecule combination. The fine-tuned model, therefore, can be used to predict affinities for novel proteins outside the training set and/or can be additionally fine-tuned given new experimental data.

The pre-trained and fine-tuned models enable both the generation and scoring of new candidates. For a given input molecule, the pre-trained model can be used to predict viable sub-sequence rearrangements similar to the mask prediction task. The fine-tuned model can then be used to predict the binding affinity for a newly generated molecule with a provided protein sequence. We utilize a genetic algorithm to automate rounds of molecule generation, scoring, and the selection of high scoring candidates.

The large scale of the pre-training and fine-tuning datasets necessitates the HPC resources of a leadership computing facility. Similar to natural language processing applications, the scale of the dataset (i.e., billions of molecules) enables the pre-trained model to learn generalizable features of molecule structure that are useful for affinity prediction. Fortunately, after pre-training and fine-tuning, the models can be used for inference or genetic algorithm optimization with modest resources (i.e., a single GPU). Therefore, our work provides models that can generalize to new protein targets, accelerate screening of potential candidates with limited or no additional fine-tuning, and be utilized throughout the research community for drug discovery efforts.

## 4 CURRENT STATE OF THE ART

### 4.1 Drug Discovery Pipelines

The complexity and challenges associated with the drug discovery process have motivated the development of pipelines to collect data and organize research efforts [12, 16]. The computational techniques in such pipelines are primarily organized around the generation and scoring of new molecules. For molecule generation, multiple different representations have been used (e.g., SMILES and graph) along with several ML model architectures (e.g., GAN, VAE, RNN) [1, 17–20]. In addition, manually-defined rules (e.g., add an atom, change atom type) have been used to generate new candidates starting from an initial population of molecules [21, 22]. For scoring candidates, both ML models and docking simulations have been used along with hybrid approaches [12, 16, 23]. The features used in ML models for scoring range from learned embeddings to chemical descriptors and molecular fingerprints [12].

Determining the correct metric for scoring and optimizing molecules is a key difficulty for the practical application of computational drug discovery pipelines. Cheminformatics packages, such as rdkit [24], provide standard heuristic metrics for chemical properties, including solubility [25], synthesizability [26], and quantitative estimation of drug-likeness [27]. However, these metrics are not specific for a given therapeutic target. Alternatively, a supervised ML model for scoring can provide customized optimization metrics but requires a suitable experimentally measured dataset [28, 29]. To overcome this difficulty, we here utilize a strategy from natural language processing, where a model is initially trained in an unsupervised manner before being fine-tuned to make specific predictions.

### 4.2 Transformers

Over the past few years, the field of natural language processing (NLP) has undergone a paradigm shift powered by the use of Transformer-based models [30]. Previously, the application of ML models was largely task specific, with a single model being trained in a supervised manner for each task (e.g., classification, similarity, entity recognition). However, with the introduction of Transformer models (e.g., BERT), training was split into two distinct stages. In the first stage (i.e., pre-training) the model is typically trained on a large corpus of text in an unsupervised manner. Unsupervised training was accomplished by using a mask prediction task, in which the model was trained to predict a given word based on context. In the second stage (i.e., fine-tuning), the pre-trained model is trained in a supervised manner on a relatively small labeled dataset. In this way, a single pre-trained model can be fine-tuned for any number of specific tasks. Models developed according to this two stage approach have achieved state-of-the-art results for a number of NLP tasks [30–32].

Advances in NLP can be directly applied to drug discovery efforts, as proteins and molecules can be represented as sequences of text. Recent efforts have indeed trained Transformer models using molecules in SMILES format for chemical property prediction tasks [2, 33–35]. Most previous work, however, has focused on a language model vocabulary of individual characters or atoms within a sequence, limiting the ability of the model to concisely represent commonly occurring chemical structures. Furthermore, the largest dataset used for pre-training contained approximately one billion molecules [2], with most investigations using fewer than 100 million [33–35]. With the success of previous Transformer models using SMILES and text data, we were motivated to increase the number of pre-training samples by an order of magnitude and utilize different model vocabularies.

Transformer models have also been trained using protein sequence data. A recent study investigated the performance of multiple model architectures on protein prediction tasks [15]. Furthermore, the outputs of pre-trained models for both molecule data and protein data can be used as embeddings for additional downstream tasks [36]. In the context of drug discovery, this enables the pretrained models to be fine-tuned on a dataset consisting of many different protein and ligand combinations with experimentally determined binding affinity. Notably, Transformer-based approaches have shown significant performance improvements for affinity prediction over alternative architectures, such as convolutional neural networks [36, 37]. In the current work, we leverage the Transformer architecture to develop a fine-tuned model capable of predicting binding affinity for novel protein targets.

### 4.3 Deep Learning at Scale

Using increasingly large training datasets poses a substantial challenge in terms of time-to-solution. Data parallelism enables many deep learning models to be trained efficiently at the scale of current supercomputers [38, 39]. Transformers are one such model; for example, large scale data parallelism has been used to dramatically reduce BERT pre-training times [40–42]. Larger Transformer models, such as Megatron-LM, have achieved performance of over 500 petaflops in mixed precision on Nvidia’s Selene supercomputer [43]. Inasmuch as a previous effort to train a BERT model using one billion molecules required approximately 4 days for pre-training [2], the potential advantage of using large scale data parallelism to enable pre-training on larger datasets with faster turnaround times is clear.

However, extreme scale data parallelism necessarily leads to extremely large batch sizes and large batch sizes can lead to instability during training which degrades model evaluation performance. While this problem is general [44], it has also specifically been observed as a scaling bottleneck in the context of developing deep learning models for drug discovery: a recent study found that an overall batch size above 4096 (across 8 or 16 GPUs) was detrimental to model training on molecule data for a variational autoencoder [1]. The recently developed LAMB optimizer has been shown to address this problem for batch sizes of up to 96 thousand, maintaining similar evaluation performance as batch size increased [40, 41].

### 4.4 Genetic Algorithms

Inspired by the mutation and selection observed in natural systems, genetic algorithms provide a useful framework for solving optimization problems across scientific and engineering disciplines [45–48]. Specifically, for drug discovery, genetic algorithms have been used in several studies to search chemical space. For example, Virshup et al. proposed a set of hand-crafted rules for mutation and recombination (e.g., add an atom, modify an atom type) to generate new compounds. The generated compounds were then selected based on diversity criteria to expand to unexplored regions of chemical space [21]. Additional studies have used genetic algorithms to optimize for drug-specific metrics (e.g., solubility and quantitative estimation of drug-likeness) [22, 46, 49, 50]. Typically, mutation operators are manually defined based on the application and not learned from the data. However, comparisons with alternative ML optimization techniques have shown that genetic algorithms perform well across a range of drug discovery tasks [50].

As an alternative to the manually defined mutation and recombination operators, molecule rearrangements can be determined by a ML model. Generative models, such as generative adversarial networks (GANs) can be used to produce molecules with desired properties [17, 18]. Furthermore, masked language models provide a useful modeling framework in which to learn viable rearrangements of molecular sequences. During pre-training the language model learns to predict missing sequences based on context. The predictions provide a ranked list of all possible substitutions for a given sub-sequence. Therefore, by sampling from the predictions, a set of mutations can be generated for a molecule without the need for manually defined rules. A similar procedure has been used to find adversarial examples for NLP applications [51, 52]. In this work, we utilize a masked language model to generate candidate molecules and then apply selection based on scoring from the fine-tuned model for binding affinity.

## 5 INNOVATIONS REALIZED

Our strategy for accelerating computational drug discovery is summarized in Figure 1. We begin by constructing the largest molecule dataset to date for pre-training a masked language model. Pretraining is performed at scale using a batch size of over a million molecules for two different tokenization schemes. We then fine-tune the language model on a dataset with binding affinities for thousands of protein targets. After developing the general pre-trained and fine-tuned models, we search for drug candidates that optimize the predicted binding affinities for a given protein.

**Figure 1:**
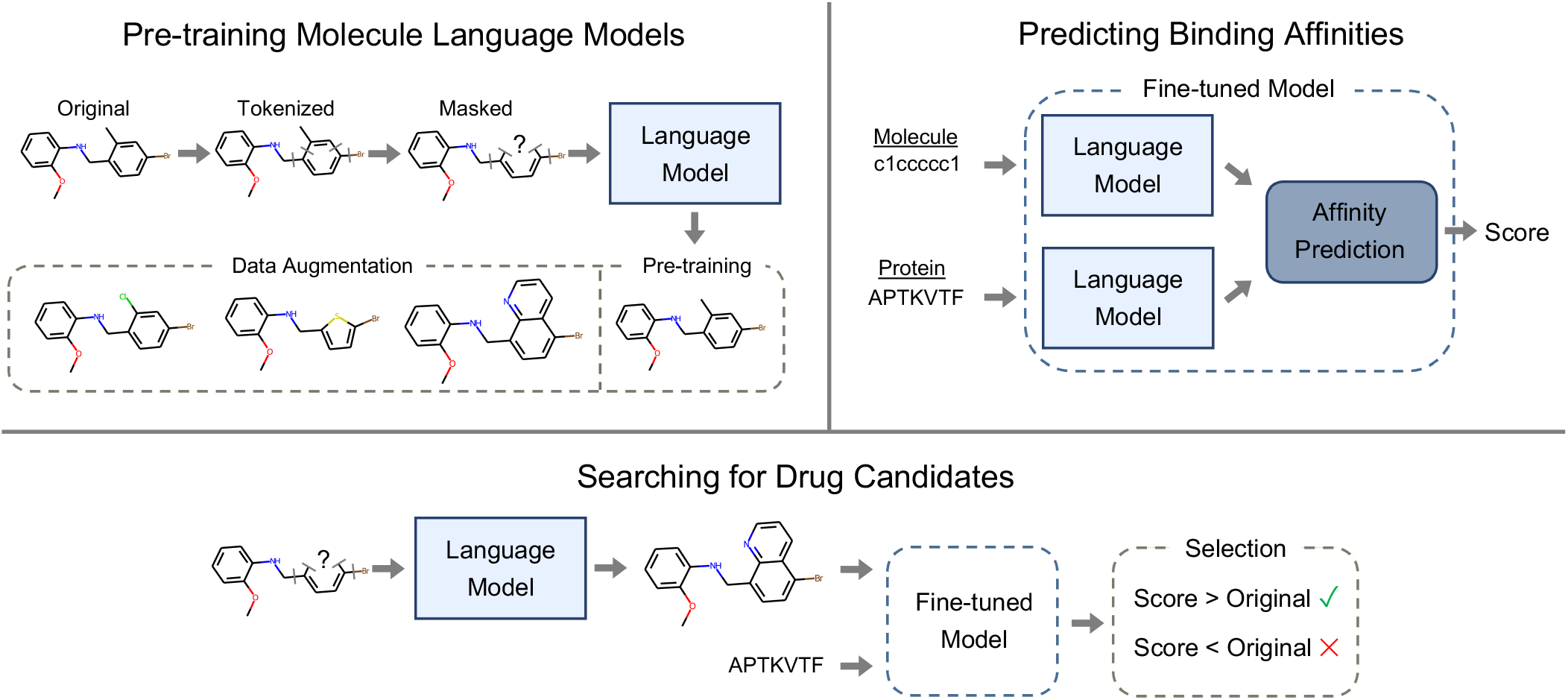
Our strategy for developing general models for drug discovery contains three components. The molecule language model is trained in an unsupervised manner; for pre-training, the model learns to reconstruct an original input molecule after masking. For data augmentation, the model predictions can be used to generate new molecules. To predict binding affinities, we use a dataset with measurements for protein and ligands to fine-tune the pre-trained model for predictions. To search for drug candidates, we use both the pre-trained language model and the fine-tuned model. The pre-trained model generates new molecules, which are scored by the fine-tuned model. Molecules are then selected based on the predicted score. The process is iterated in the search for optimized drug candidates.

### 5.1 Pre-training Molecule Language Models

#### 5.1.1 Dataset Generation

Motivated by the success of Transformer models (i.e., BERT) for a range of natural language processing tasks, recent efforts have investigated using the SMILES text representation of molecules to train a masked language model [2, 33–35]. Large compound libraries such as Enamine *REAL* database [14] can be used for the unsupervised pre-training stage, before the model is fine-tuned for a desired prediction task. Although compound libraries provide a valuable source of training data, the overwhelming size of chemical space ensures that many potentially useful compounds will be excluded from current collections. Current state-of-the-art generative and masked language models have reached a training data size of approximately 1.1-1.6 billion compounds, pushing the boundaries of currently available compound libraries and compute resources [1, 2]. As a step toward enabling larger explorations of chemical space, here we utilize the Enamine *REAL* database as a starting point to generate a training dataset with ∼9.6 billion unique molecules.

Our strategy for dataset augmentation is motivated by the pretraining stage for masked language models. During pre-training, random sequences of an input molecule SMILES are masked, and the model is trained to predict the identity of the masked sequences based on the surrounding context. We, therefore, use a pre-trained model, developed using Enamine as the training data, to predict possible structural rearrangements for a given molecule as shown in Figure 1. We also use the pre-trained model to combine two molecules; initial sequences are chosen from each respective molecule, and a mask is placed in between. To be included in the training set, all produced molecules must be valid [24] and have a normalized synthesizability [18, 26] score above 0.3. To arrive at the final dataset of 9.57 billion molecules, we applied random structural rearrangements to molecules in the Enamine dataset with a maximum of five masks per molecule. The top 5 predicted rearrangements from the pre-trained model were considered, resulting in the dataset growing from approximately 1.34 billion unique compounds to 4.14 billion unique compounds. Another round of rearrangements for the expanded dataset was accompanied by combinations of molecules to generate the final training set. All molecules were converted to canonical form using rdkit [24] and only unique molecules were retained in the final dataset. As shown in Figure 2, the histograms for drug-likeness, normalized synthesizability, and solubility did not substantially change between the original data and the augmented data, although the total number of compounds substantially increased.

**Figure 2:**
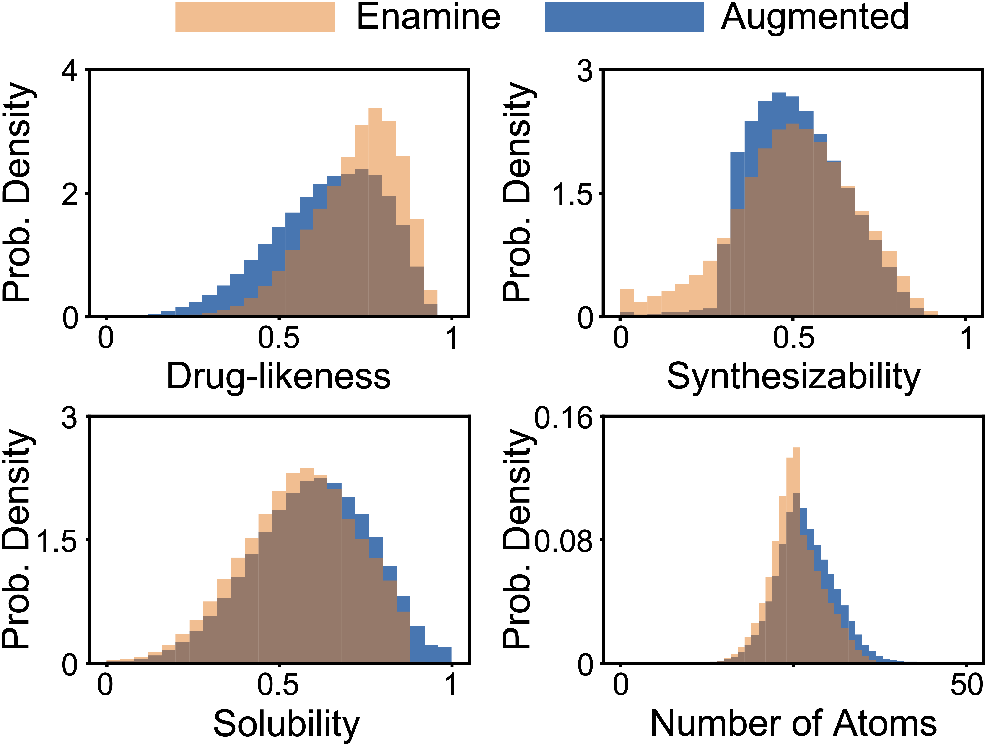
Although the augmented dataset contains roughly 7 times more molecules than the original dataset, the histograms show that the augmentation strategy largely preserves the distribution of multiple molecule metrics. For Synthesizability, generated molecules for data augmentation were required to have a score above 0.3, resulting in the observed sharp decline in the histogram. For drug-likeness, no constraints were placed on the augmented data, which resulted in a decrease in typical scores relative to Enamine.

#### 5.1.2 Tokenization

The process of tokenization is used to convert any given sequence of text into a format that can be recognized by the model. This is accomplished by constructing a vocabulary for the model, which consists of a mapping between sub-sequences of text and unique integer ids. One common method, WordPiece tokenization [53, 54], builds the model vocabulary by assembling all unique single characters and then including commonly occurring sequences of increasing length. Although a collection of different tokenization methods have been used for NLP tasks, for molecule language models a simple vocabulary based on single atoms and characters has predominantly been used [2, 19, 20, 33, 35, 55, 56].

Here, we utilized two different tokenization methods, the standard single atom and character Regex [56] and WordPiece tokenization. As shown in Figure 3, the vocabulary generated by WordPiece tokenization enables large sub-sequences of a SMILES string (e.g., a benzene ring) to be represented as a single token. Notice that for the Regex tokenizer, the vocabulary will contain only individual characters and atoms, so commonly occurring chemical structures cannot be assigned to a single token. Also, the size of the vocabularies for the two tokenizers is drastically different, with the WordPiece tokenizer having 3·10^4^ different tokens, while the Regex tokenizer has around 200. Given the substantially different representations of molecules produced by the two tokenizers, we utilized both methods for pre-training and fine-tuning tasks to determine the impact of tokenization on fine-tuning and molecule generation performance.

**Figure 3:**
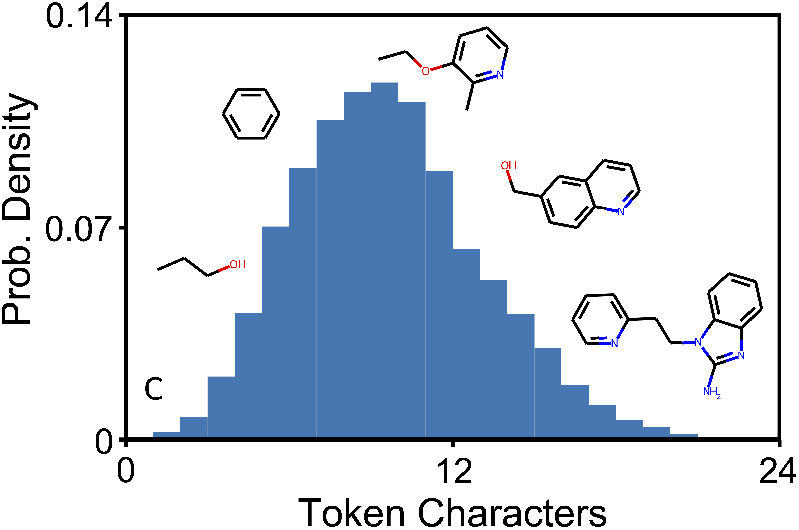
The vocabulary generated by WordPiece tokenization represents commonly occurring sub-sequences from the training data as individual tokens. The histogram shows the distribution of number of characters for all tokens in the vocab along with the chemical structure for sample tokens of different length.

#### 5.1.3 Pre-training with Large Batch Sizes

We trained BERT models using the LAMB optimizer with both the Regex and WordPiece tokenizers utilizing different numbers of nodes on Summit. Each node contained 6 GPUs, each having a single partition of 3.95 10^5^ unique molecules. The batch size per GPU was set to 240 (80 with 3 gradient accumulation steps); therefore, the total batch size is given by 1440 (i.e., 240·6) multiplied by the number of nodes [57]. At 1000 nodes, this results in over 1.4 million molecules per batch. As shown in Tables 1-2, even with large batch sizes, the model can be trained successfully, as evidenced by the validation accuracy for mask prediction. Validation accuracy was determined by evaluating the model on a hold-out set of 10^5^ molecules; for each molecule a random number of masks (up to 5 for Regex and up to 3 for WordPiece) were sampled and used to replace tokens. Each pre-training run consisted of 7 epochs, with model checkpoints saved and validation accuracy determined after each epoch. The maximum accuracy across checkpoints is shown. Notice that a comparison of accuracy between Table 1 and Table 2 should not be made, as the mask prediction task is substantially easier for the Regex tokenizer (i.e., only single atoms or characters are predicted).

**Table 1:** Validation Accuracy for Pre-training Runs with WordPiece Tokenizer.

| Nodes | Molecules | Batch Size | Accuracy |
| --- | --- | --- | --- |
| 1 | $2.4 \cdot 10^6$ | $1.4 \cdot 10^3$ | 0.760 |
| 10 | $2.4 \cdot 10^7$ | $1.4 \cdot 10^4$ | 0.783 |
| 100 | $2.4 \cdot 10^8$ | $1.4 \cdot 10^5$ | 0.797 |
| 1000 | $2.4 \cdot 10^9$ | $1.4 \cdot 10^6$ | 0.798 |
| 1008 | $9.6 \cdot 10^9$ | $1.5 \cdot 10^6$ | **0.808** |
| 4032 | $9.6 \cdot 10^9$ | $5.8 \cdot 10^6$ | 0.801 |

**Table 2:** Validation Accuracy for Pre-training Runs with Regex Tokenizer.

| Nodes | Molecules | Batch Size | Accuracy |
| --- | --- | --- | --- |
| 1 | $2.4 \cdot 10^6$ | $1.4 \cdot 10^3$ | 0.845 |
| 10 | $2.4 \cdot 10^7$ | $1.4 \cdot 10^4$ | 0.866 |
| 100 | $2.4 \cdot 10^8$ | $1.4 \cdot 10^5$ | 0.878 |
| 1000 | $2.4 \cdot 10^9$ | $1.4 \cdot 10^6$ | 0.882 |
| 1008 | $9.6 \cdot 10^9$ | $1.5 \cdot 10^6$ | **0.889** |

For the full dataset, we used two different training configurations. First, we used 1008 nodes, with 4 partitions of 3.95·10^5^ unique molecules per GPU. For comparison, we also performed pre-training on over 4000 nodes for the WordPiece tokenizer with a single partition per GPU (last row of Table 1). Both pre-training runs used a warmup of 1 epoch. As expected from previous studies [41], the increased batch size for the 4032 node run resulted in decreased validation accuracy; however, it is notable that a batch size of nearly 6 million incurred only a slight decrease in performance, suggesting that distributed training for even larger molecule datasets is possible. The 1008 node runs were completed in 8 hours each for 7 epochs; the 4032 node run was stopped after failing to increase validation accuracy (maximum was at 5 epochs), taking less than 2 hours. For downstream tasks, such as fine-tuning, we used the models trained on 1008 nodes for the WordPiece and Regex tokenizers.

### 5.2 Predicting Binding Affinities

To determine whether a given drug molecule binds to a target molecule, i.e., a protein, both the candidate molecule and the amino acid sequence of the protein need to be embedded. Then, to predict the binding affinity, hidden layers are added that accept the concatentation of the two embeddings as inputs. The predictive power of such an ensemble model is chiefly determined by the expressive power of the individual embeddings. Therefore, we expect that using powerful pre-trained models to embed the molecules results in superior performance on the downstream task. Here, we focus on regression to predict the numerical value of the binding affinity, however, the model architecture lends itself equally well to classification. Pre-trained embeddings and extra layers are fine-tuned simultaneously, i.e., with all weights adjustable, on a labeled data set of 1.67·10^6^ receptor amino acid sequences and ligand SMILES, with binding affinities. Note that this dataset used for fine-tuning is much smaller than the datasets used for pre-training (Tables 1&2).

#### 5.2.1 Binding Affinity Dataset

We curated a dataset [58] of binding affinities by concatenating the entries of the BindingDB [59], PDBBind-cn [60], BindingMOAD [61], and BioLIP [62] databases, following the example of Ref. [63]. Records containing *K*_*i*_, 1/*K*_*a*_, *K*_*b*_ and IC_50_ values were retained and converted to pKd= −log_10_ *K*_*d*_ [M] units, and MACCS fingerprints were calculated on the molecules to remove duplicates, resulting in 1,670,637 protein sequences, SMILES strings and binding affinities.

#### 5.2.2 Model Architecture

Figure 4 shows the neural network architecture used to predict affinities. For embedding molecules, we use the tokenizers and pre-trained models discussed above. For embedding proteins, we make use of the readily available pre-trained ProtBERT model [64], where every token is a letter in the amino acid alphabet. The embeddings are fed to a cross-attention module [65]. The purpose of the cross-attention layer is that molecule subunits attend to amino-acids in the protein sequence, and *vice versa*. This architecture represents the physical situation in which the molecule makes well-defined atom-atom contacts with the protein. However, the model is not constrained to learn real physical contacts, and importantly, it is not given any information about which residues belong to the active site of the protein, which it has to learn by itself from the given correlation between structures and binding affinities. Despite the physical motivation behind its architecture, the model is still to be considered as a ‘black-box’. It cannot be expected that the cross-attention weights directly correspond to observable physical contacts, as sometimes suggested [63, 65].

**Figure 4:**
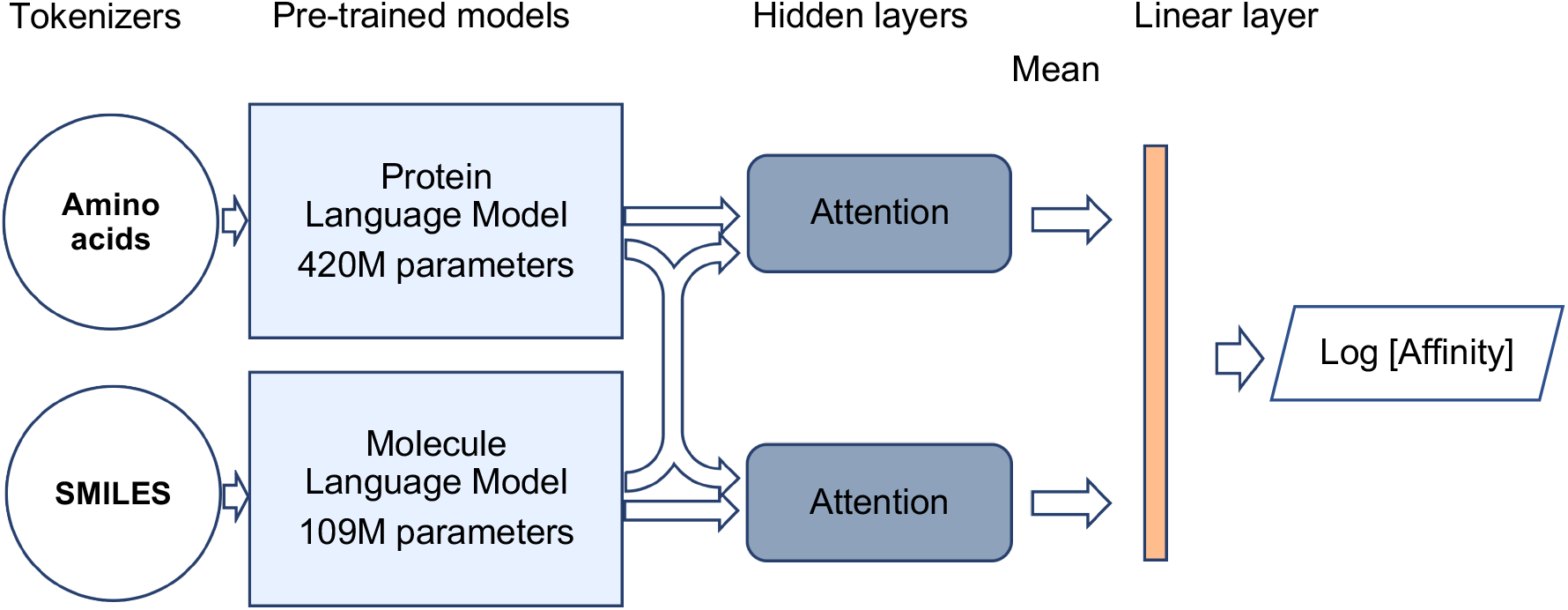
Architecture of the affinity prediction model used for fine-tuning. It uses two independent models for protein sequences (Protein Language Model) and molecule SMILES (Molecule Language Model) connected to a cross-Attention module that predicts the logarithm of the binding affinity.

The hidden layer outputs of the cross-attention module are concatenated, their mean is taken over the sequence length and they are connected to a linear layer to predict the binding affinity. The model is fine-tuned by minimizing the mean-squared error (MSE) between the predicted and the experimental affinity. We validated the model on a hold-out set from the training data as well as three additional datasets as shown in Tables 3-4. We characterize the ability of the model to correctly reproduce the order of the experimental affinity values by the Spearman-*ρ* rank correlation coefficient (higher is better) [69], as well as the mean-squared error for the predicted affinity (lower is better). We calculate the uncertainty in the reported values using the bootstrap method with *n* = 500 samples. Notably, the Regex tokenizer (and associated ensemble) outperforms the WordPiece tokenizer for certain datasets (i.e., Hold-out and Kinases), but underperforms for others (i.e., PLpro), suggesting that different molecule representations may be suitable for different affinity prediction tasks.

**Table 3:** Validation of the affinity prediction on different test sets, with Regex tokenizer for SMILES. Shown are Spearman *ρ* rank correlation coefficient, and mean-squared error (MSE) for the ensemble and the individual models. Values in parentheses indicate the uncertainty of the last reported digit.

| Test set | Ensemble |  | Model 1 |  | Model 2 |  | Model 3 |  |
| --- | --- | --- | --- | --- | --- | --- | --- | --- |
| | $\rho$ | MSE | $\rho$ | MSE | $\rho$ | MSE | $\rho$ | MSE |
| Hold-out | **0.881**(2) | **0.69**(1) | 0.866(3) | 0.77(1) | 0.865(3) | 0.79(1) | 0.866(3) | 0.75(1) |
| Kinases [66] | **0.38**(1) | **1.67**(3) | 0.35(1) | 1.80(3) | 0.35(1) | 1.88(3) | 0.35(1) | 1.80(3) |
| Mpro [67] | **0.39**(2) | 1.82(5) | 0.28(2) | **1.79**(5) | 0.37(2) | 2.24(6) | 0.32(2) | 1.82(5) |
| PLpro [68] | 0.54(8) | 0.8(1) | **0.55**(8) | 0.9(1) | 0.25(10) | 0.67(11) | 0.45(8) | 1.3(2) |

**Table 4:** Validation of the affinity prediction on different test sets, with WordPiece tokenizer for SMILES. Shown are Spearman *ρ* rank correlation coefficient, and mean-squared error (MSE) for the ensemble and the individual models. Values in parentheses indicate the uncertainty of the last reported digit.

| Test set | Ensemble |  | Model 1 |  | Model 2 |  | Model 3 |  |
| --- | --- | --- | --- | --- | --- | --- | --- | --- |
| | $\rho$ | MSE | $\rho$ | MSE | $\rho$ | MSE | $\rho$ | MSE |
| Hold-out | **0.864**(3) | **0.72**(1) | 0.830(3) | 0.94(2) | 0.846(3) | 0.84(1) | 0.854(3) | 0.77(1) |
| Kinases [66] | **0.30**(1) | **2.03**(3) | 0.23(1) | 2.90(4) | 0.26(1) | 1.95(3) | 0.30(1) | 1.94(3) |
| Mpro [67] | **0.37**(2) | 1.23(5) | 0.33(3) | **1.23**(5) | 0.29(2) | 1.49(5) | 0.31(3) | 1.44(5) |
| PLpro [68] | 0.57(8) | 0.71(10) | 0.51(8) | **0.45**(7) | **0.61**(7) | 0.76(10) | 0.30(10) | 1.7(2) |

Figure 5 demonstrates the performance for the validated and transferable model on a binary classification task, using the metrics of precision and recall. Here, we impose a threshold of 5 *μ*M (Mpro) and 1 *μ*M (PLpro) on the experimental IC50 value (lower is better) to label active molecules. These thresholds are typical of more potent non-covalent inhibitors for the Mpro and PLpro targets. We use sampling from the normal distribution of affinities implied by the mean and the variance of the ensemble model to estimate the confidence intervals, as well as the standard error from *n* = 500 bootstrap samples. Remarkably, the model achieves a maximum precision of 0.60 both for Mpro and PLpro, meaning that 60% of the highest scoring molecules are true actives, which suggests excellent virtual screening performance.

**Figure 5:**
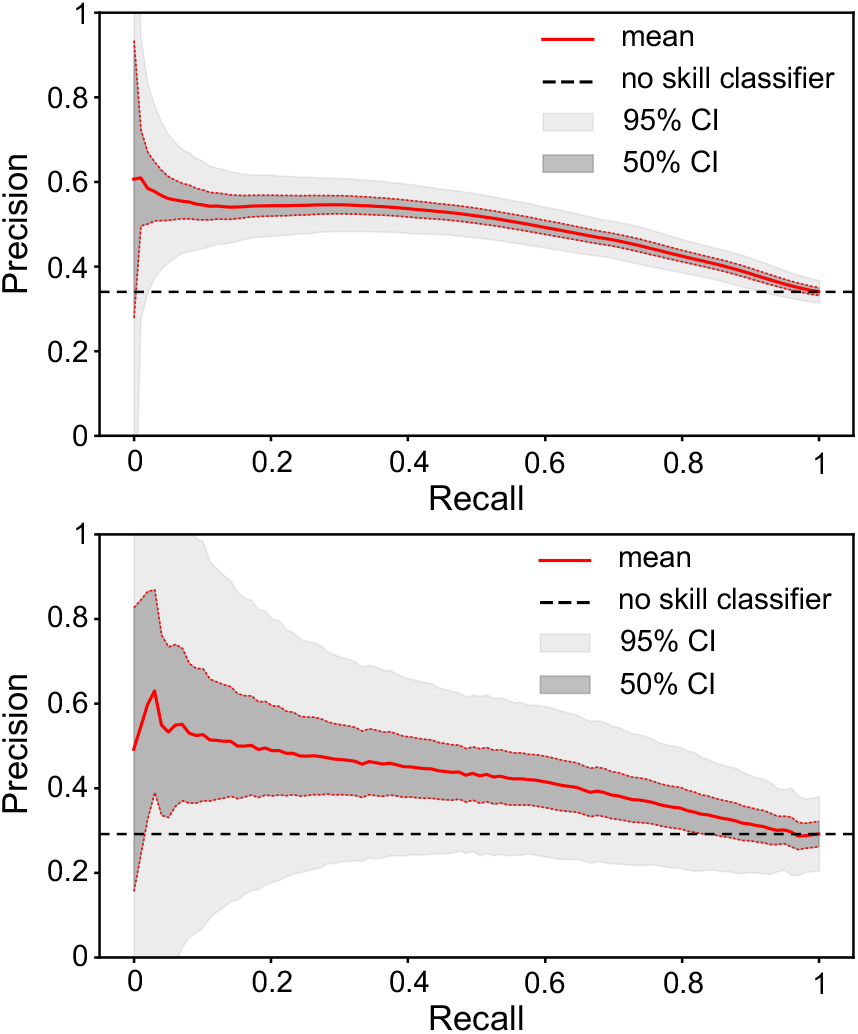
Transferability and virtual-screening performance of the fine-tuned model on two experimental SARS CoV-2 protein affinity data sets, Mpro (top, from Ref. [67]) and PLpro (bottom, from Ref. [68]), showing precision [=*tp*/(*tp* + *f p*)] as a function of recall [=*tp*/(*tp* + *f n*)].

### 5.3 Searching for Drug Candidates

Similar to the strategy we used for data augmentation, the pretrained model can easily be adapted to generate rearrangements for a given population of molecules. Here we utilize three different types of rearrangements for a given tokenized molecule: insertion, deletion, and replacement. For insertion, a mask is randomly inserted in between existing tokens or at the beginning or end. For deletion, a mask randomly replaces two adjacent existing tokens. With replacement, a single existing token is randomly masked. To search for new drug candidates, we randomly sampled up to 5 masks and a rearrangement type for molecules in the population. In addition, we consider recombination by randomly sampling two molecules, selecting a sub-sequence from each and inserting a mask in between. A canonical and randomized SMILES were used to represent each molecule before masking, and the top 10 molecules predicted by the pre-trained model were used as candidates. As a starting population, we used molecules from the validation set for pre-training. Only unique molecules were retained in the population, as determined by canonical SMILES computed using rdkit [24].

To score the molecules generated through rearrangements, we utilized three metrics: normalized synthesizability, quantitative estimation of drug-likeness, and the affinity predictions of the finetuned model. The predicted score for the affinity was divided by 10 and clipped between 0 and 1 to generate a normalized affinity metric. The harmonic mean of the three metrics was then used to define the fitness of a given candidate. To find optimized candidates, an initial population of 10^4^ molecules was used from the validation set for pre-training. Then, masked rearrangements were applied to 5·10^3^ samples and recombination was applied to 5·10^3^ sampled pairs. The resulting molecules were added to the population and ranked according to fitness; the top 10^4^ overall were retained as the starting population for the next generation. Based on the Hold-out fine-tuning results, we selected Model 3 with a Regex Tokenizer to predict scores. For molecule rearrangements, we used the pretrained model with WordPiece Tokenizer as it generated higher fitness scores than the corresponding Regex Tokenizer.

As shown in Figure 6 (top two rows), 50 generations of optimization to search for Mpro inhibitors resulted in a substantial shift in the distributions of the three optimized metrics. Notably, the mean for the affinity score increases, with the maximum generated molecule having a normalized affinity score greater than 0.9. By optimizing for synthesizability and drug-likeness in addition to affinity, the generated molecules are constrained by useful heuristic scoring functions for drug discovery [26, 27]. We also show the top scoring molecule in the final population for each respective metric. The three examples show that optimization successfully found molecules with higher predicted affinities while maintaining high synthesizability and drug-likeness scores. The bottom two rows of Figure 6 show the corresponding results for PLpro. The only change to the genetic algorithm is the input protein sequence, highlighting the generality of our approach.

**Figure 6:**
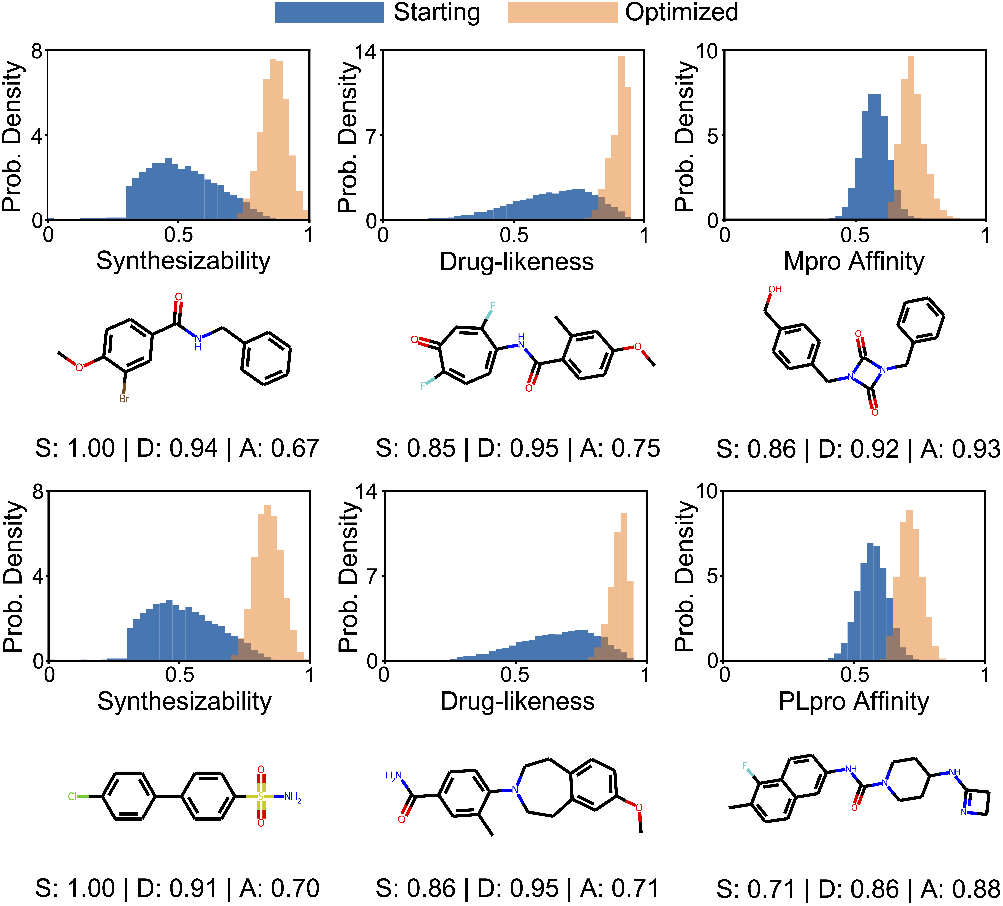
Results from optimizing molecules for the harmonic mean of synthesizability, drug-likeness, and affinity. The top two rows show results for Mpro; the bottom two rows show results for PLpro. The histograms show the changes in the probability distributions from the starting population to the optimized population after 50 generations. The three examples show the highest scoring molecules for each respective metric in the optimized population

## 6 HOW PERFORMANCE WAS MEASURED

### 6.1 Applications Used to Measure Performance

In this study, we performed pre-training, fine-tuning, and a genetic algorithm on Transformer models. All models are written in PyTorch using the Hugging Face Transformers API [70]. Models are pre-trained and fine-tuned with DeepSpeed [42], a high performance wrapper for distributed Transformer training. Model training is performed with data parallelism using DeepSpeed’s fused-kernel LAMB optimizers. Sharded I/O is performed using the WebDataset library [71]. As pre-training of the molecule language model is by far the most computationally expensive stage of the study, it is the focus of the performance analysis.

The architecture used for the molecule language model is BERT-base, which has approximately 109 million learnable parameters. Pre-training of the model is performed with data parallelism, in which each GPU trains the model on separate data. Communication takes the form of a global asynchronous AllReduce which is performed during backpropagation on each batch.

### 6.2 Measuring Performance

Performance of molecule language model pre-training was measured in two respects. First, sustained performance was measured using built-in timers which report the total wall clock time elapsed during training and the time for I/O operations (specifically, saving checkpoints and trained models). Additionally, to measure peak application performance relative to theoretical machine peak, mixed precision floating point operations per second (FLOPs) are computed using the DeepSpeed FLOPS Profiler.

### 6.3 System

#### 6.3.1 Hardware

Performance was measured on the Summit supercomputer at the Oak Ridge Leadership Computing Facility at ORNL [72]. Summit is comprised of 4674 IBM Power System AC922 nodes which are arranged in a non-blocking Fat Tree topology with dual-rail EDR InfiniBand interconnect. Each node has two IBM Power9 CPUs, six Nvidia 16 GB V100 GPUs, and 512GB of main memory. The V100 device has an estimated peak performance of 14 teraflops for single precision (FP32) and 112 teraflops for mixed precision using the Tensor Cores, which are capable of performing matrix multiply in FP16 with FP32 accumulation for some kernels. Consequently, Summit’s peak performance for mixed precision is approximately 3.1 exaflops.

#### 6.3.2 Software

Summit runs the Red Hat Enterprise Linux 8 operating system and uses the IBM LSF job scheduler. Our Python-based software stack uses Open Cognitive Environment v1.2.0, PyTorch v1.7.1, Transformers v4.5.1, DeepSpeed v0.4.5, and WebDataset v0.1.62. The GPU libraries include CUDA 11.0.3, NCCL 2.7.8, and cuDNN 8.0.4.

## 7 PERFORMANCE RESULTS

### 7.1 Node Level Optimization

The results of node level optimization for molecule language model pre-training are shown in Figure 7, in terms of runtime per epoch plotted versus a series of successive optimizations. In this setting, data parallelism is applied to train the molecule language model across the six GPUs of a Summit node. The baseline case uses the Adam optimizer, single precision arithmetic, and the largest batch size per device (96) for which the model fits in GPU memory. Enabling mixed precision with the V100 Tensor Cores decreases runtime by approximately 43%. In addition, mixed precision enables a larger per device batch size (128) to be used while keeping the model within GPU memory, further decreasing runtime per epoch by about 15%. While the decision to switch to the LAMB optimizer was motivated by large batch sizes, as discussed in section 5.1.3, DeepSpeed’s fused LAMB optimizer implementation also improves performance by roughly 15%. Finally, leveraging LAMB’s stability for very large batches, we changed the batch size to 80 and added 3 gradient accumulation steps, for an effective batch size per device of 240. While the addition of gradient accumulation does not improve node level performance, this configuration enables better scaling at very high node counts due to reduced communication frequency and, after hyperparameter optimization, maintains comparable accuracy.

**Figure 7:**
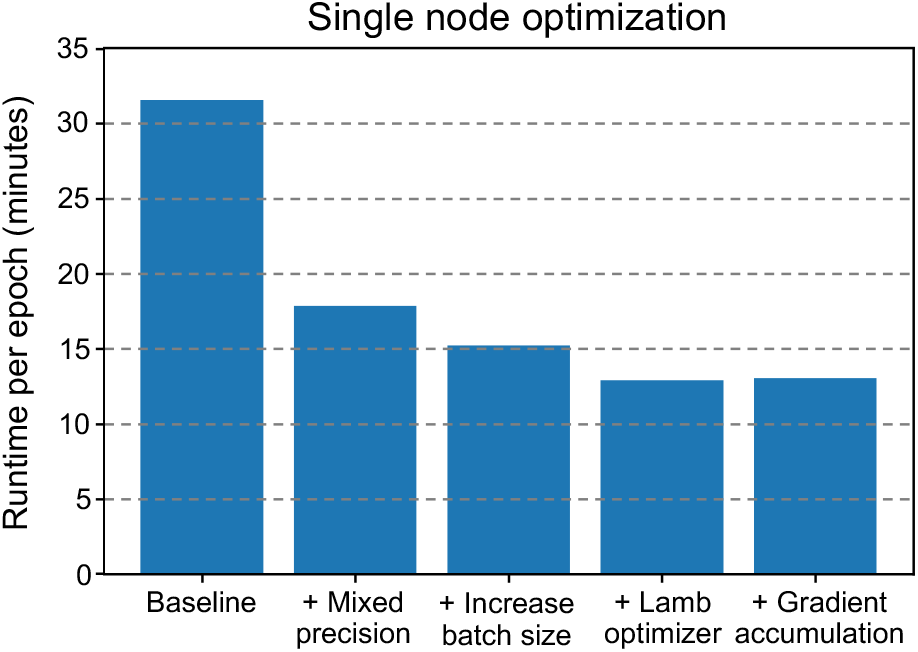
Single node algorithmic and performance optimization for pre-training the molecule language model: (1) base-line with Adam optimizer, (2) use of mixed precision arithmetic on V100 Tensor Cores, (3) larger per device batch size enabled by mixed precision, (4) fused LAMB optimizer, and (5) optimal balance of batch size per device and gradient accumulation steps for single node performance and scalability.

### 7.2 Scaling

As this study incorporates the largest molecule dataset ever used for pre-training, the primary focus for scalability was weak scaling.

In Figure 8, we assess the weak scaling of pre-training the molecule language model on Summit. For weak scaling, the problem size per device is kept constant at 3.95 10^5^ molecules. The training configuration is that identified in section 7.1, extended to the multi-node setting, with data parallelism used to train a single model across the given number of nodes. These runs are the production runs in section 5.1.3 with the WordPiece tokenizer, and therefore include I/O operations to save checkpoints and the final trained model. Parallel efficiency for weak scaling from 1 to 4032 Summit nodes is measured at 68.0%. However, a significant amount of performance degradation at large node counts is due to I/O; when I/O time is subtracted out, parallel efficiency over the same interval improves to 83.3%. In combination with the validation accuracy results from Tables 1 and 2, this clearly indicates that pre-training can be extended to incorporate unprecedented molecule dataset and batch sizes without significantly compromising computational efficiency or accuracy.

**Figure 8:**
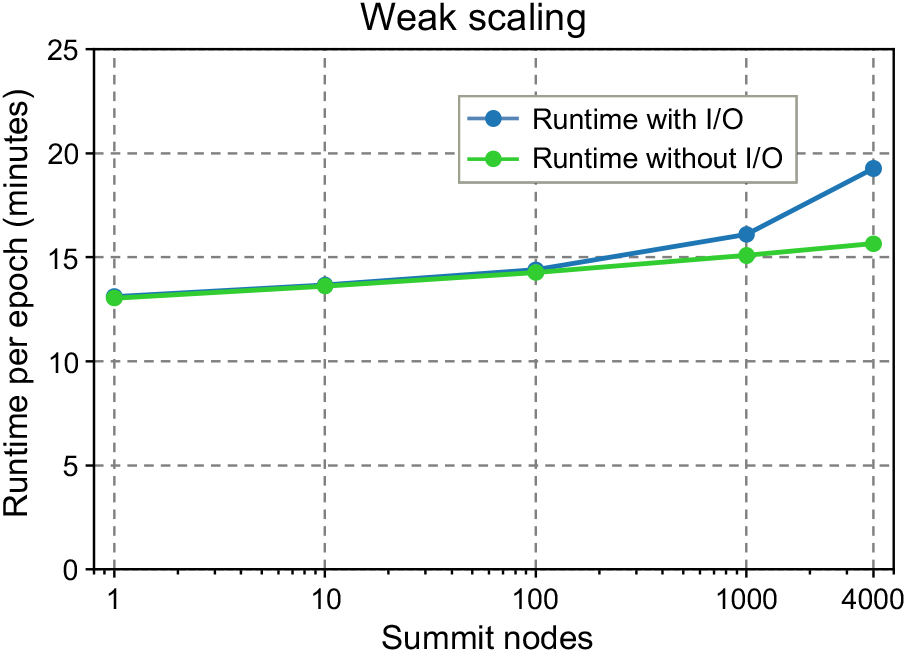
Weak scaling of molecule language model pretraining on Summit for a constant 395 thousand molecule problem size per GPU. I/O operations are saving checkpoints and trained models.

Strong scaling of the molecule language model pre-training is shown in Figure 9 from 25 to 1600 Summit nodes. The total problem size is kept constant at approximately 1.9 billion molecules, with the same training configuration as for weak scaling. However, as full runs could not be completed within facility wallclock limits for many node counts, the reported runtime per epoch is measured for the first 0.25 epochs for the 25 node job and for the first epoch for all other node counts. Strong scaling is near linear from 25 to 400 Summit nodes and maintains approximately 67.4% parallel efficiency at 1600 nodes versus the 25 node baseline.

**Figure 9:**
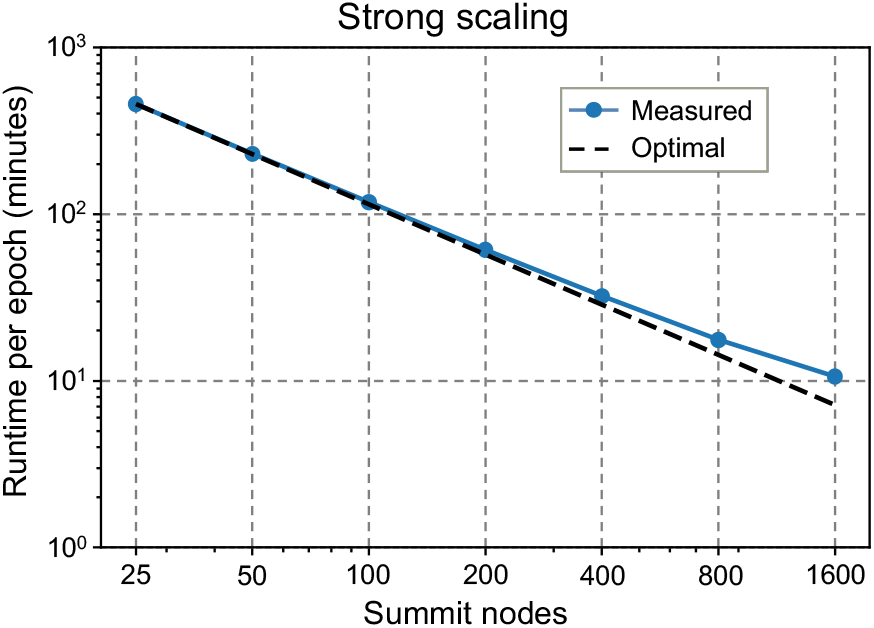
Strong scaling of molecule language model pretraining on Summit for a constant total problem size of ∼1.9 billion molecules.

### 7.3 Peak Performance

Table 5 shows the peak performance achieved for molecule language model pre-training on Summit during the largest weak scaling run from section 7.2. On 4032 nodes, peak performance of approximately 603.4 petaflops in mixed precision is achieved. As Summit’s theoretical peak at this node count is projected at∼ 2.71 exaflops, our result represents about 22.3% of this mixed precision peak.

**Table 5:** Peak performance for molecule language model pre-training on Summit in mixed precision floating point operations per second (FLOPs).

| Nodes | Molecules | Batch Size | FLOPs |
| --- | --- | --- | --- |
| 4032 | $9.6 \cdot 10^9$ | $5.8 \cdot 10^6$ | 603.4 petaflops |

## 8 IMPLICATIONS

### 8.1 ML Models for Drug Discovery

Supervised training of ML models for drug discovery poses a key difficulty in terms of both time and resources, as a labeled dataset must be created for each new therapeutic target. As demonstrated by language models for both text [30] and chemical sequence data [2], an alternative approach is to leverage large unlabeled datasets to train a general model. The general model is then fine-tuned on a relatively small dataset for a specific task of interest. Although finetuning still requires supervised training, unsupervised pre-training has enabled state-of-the-art results across a range of tasks with limited labeled data [30, 32]. Here, we have taken the shift towards general models a step further by using pre-training and fine-tuning tasks that can generalize to any protein and molecule sequence.

The ability to pre-train molecule language models with large batch sizes enables an unprecedented exploration of chemical space. Current chemical databases provide hundreds of billions of molecules, but contain only a small fraction of potentially synthesizable molecules [73]. Through the process of tokenization and mask prediction, language models can leverage large datasets to automatically learn commonly occurring subsequences (i.e., structural components) and possible rearrangements for effective searches of chemical space.

By combining a pre-trained model for molecule and protein sequences, the fine-tuning task can leverage data from many previous experimental investigations. Furthermore, additional modeling techniques, such as docking simulations, could be used to augment the training data in novel regions of interest. Recent development in protein structure prediction [74] leverage both sequence and spatial information to increase predictive performance. Although there is still much work to be done to make a truly general model for drug discovery, the increase in unsupervised and semi-supervised approaches to training along with the increase in available experimental and simulation data for protein and ligand interactions makes possible the development of off-the-shelf models that generalize across therapeutic targets. Developing a generalizable model is key to reducing the time for discovering and screening new targets in an emerging pandemic, such as COVID-19.

Using a genetic algorithm coupled with a pre-trained language model for optimization enables incremental exploration and refinement from known drugs as well as population searches. For example, a certain subsequence of a known compound can be masked, and the language model can predict the most likely rearrangements. Heuristic metrics and ML models can then be used to analyze the expected impact of structural changes. The intuitive process of making incremental changes during exploration can be used to complement and guide researcher intuition during the drug discovery process. Therefore, our suggested optimization strategy provides a natural framework for both fully-automated and user-guided exploration of chemical space.

### 8.2 HPC Resources for Model Development

The pre-training phase of developing a language model requires substantial computational resources; here, we utilized thousands of GPUs and corresponding node hours to complete training. Similarly, as the labeled dataset and the compound library for screening grows, fine-tuning and inference can also necessitate the resources of a leadership computing facility. Although these resource requirements provide an excellent use case and motivation for the continued development of HPC systems, they generate challenges for the utilization and training of models throughout the research community. Fortunately, the pre-trained models can be leveraged for inference and to some extent fine-tuning applications with only a single GPU.

For fine-tuning tasks with a specific protein target or chemical property, training can typically be done without the need for HPC resources. For inference, our results show that a genetic algorithm using the pre-trained model for mutations can be used to generate optimized candidates without the need for large scale model evaluations. Furthermore, the genetic algorithm approach provides an interpretable scheme for modifying a single molecule. The mask predictions and scores can be inspected to determine single mutations that lead to higher scores for a given metric. Also, the reported genetic algorithm runs from this work used only a single V100 GPU for less than 10 hours for optimization. Therefore, although the pre-trained models require substantial computational resources for training, the models can be used for exploration throughout the research community.

## 9 ACKNOWLEDGMENTS

We thank Jerry Parks for help in preparing the test dataset of PLpro inhibitors. This research used resources of the Oak Ridge Leadership Computing Facility, which is a DOE Office of Science User Facility supported under Contract DE-AC05-00OR22725. This research was supported by the Exascale Computing Project (17-SC-20-SC), a collaborative effort of the U.S. Department of Energy Office of Science and the National Nuclear Security Administration. This work was supported by DOE CARES emergency funding to the National Center for Computational Sciences at ORNL through the Advanced Scientific Computing Research (ASCR) program.

